# Alternative female and male developmental trajectories in the dynamic balance of human visual perception

**DOI:** 10.1101/2021.02.11.430816

**Authors:** Gergő Ziman, Stepan Aleshin, Zsolt Unoka, Jochen Braun, Ilona Kovács

## Abstract

The numerous multistable phenomena in vision, hearing and touch attest that the inner workings of perception are prone to instability. We investigated a visual example – binocular rivalry – with an accurate no-report paradigm, and uncovered developmental and maturational lifespan trajectories that were specific for age and sex. To interpret these trajectories, we hypothesized that conflicting objectives of visual perception – such as *stability* of appearance, *sensitivity* to visual detail, and *exploration* of fundamental alternatives – change in relative importance over the lifespan. Computational modelling of our empirical results allowed us to estimate this putative development of stability, sensitivity, and exploration over the lifespan. Our results confirmed prior findings of developmental psychology and appear to quantify important aspects of neurocognitive phenotype. Additionally, we report atypical function of binocular rivalry in autism spectrum disorder and borderline personality disorder. Our computational approach offers new ways of quantifying neurocognitive phenotypes both in development and in dysfunction.

## 1 Introduction

An 80 year old grandmother and her 8 year old grandson will react very differently to the same stimulus, e.g. a loud truck passing the street outside the house. The child will be engaged by the sound, turn his body toward the sound source, perhaps even drop his toys and rush to the window for a better look. Grandma, on the other hand, may not even lift her eyes from the page she was reading. While the child cannot help being over-sensitive to novel stimuli, even when it is occupied elsewhere, the grandmother – given her extensive past sensory experience – likely remains indifferent. This scenario illustrates the main hypothesis of this paper: adaptive human functioning balances conflicting objectives, and this balance changes throughout life.

Human neurocognitive development is a multidimensional and ever-changing process determined both by biological mechanisms and the environment over the lifespan. Details on the protracted maturation of human structural brain connectivity are emerging with the advance of brain imaging technology [1–5]. It is becoming increasingly clear from such findings that lifespan trajectories of brain organization improve our current, fragmented view of human development. Nonetheless, little is known about the trajectories of active neurocognitive adaptation to the environment, in other words, about the lifetime development of our own behavioural phenotype.

Because of the aforementioned complexity of human growth, the description of the neurotypical behavioural phenotype ought to be complex as well. It does not seem meaningful to ask what the typical brain structure of an adult human is. One needs to specify at least age and sex, since the brain is continually reorganizing, with its development extending into adulthood [6, 7], and as its structure shows differences between the sexes throughout development [8–10]. Adolescence, in particular, is a period of substantial changes in brain structure –– notably, in the association cortex, connectivity is remodelled [11], while cortical shrinkage and myelination are accelerated [12, 13]. These organizational changes are coupled with a high vulnerability to mental health disorders [14, 15]. Parallel to the changes in the brain’s structural organization that take place across the lifespan, we should expect changes in the way it adapts to the environment. Further differences should be expected in the case of atypical development underlying mental health disorders, whether resulting from developmental conditions, environmental factors, or a combination of both.

The aim of the present work is to characterize lifespan trajectories for visual perception. Sensory perception is generally a promising target for such an investigation, because there is considerable flexibility in how sensory perception balances its several conflicting objectives. Accordingly, over the course of development and maturation, sensory dynamics may well have to change in order to accommodate different behavioural strategies or increasing sensory experience.

That successful interaction with volatile and unpredictable environments necessitates trades between conflicting objectives is a central insight from reinforcement learning [16, 17]. For example, relying on past experience in order to make choices in perception or action typically creates an “exploration-exploitation-dilemma” [17–19]. Here, the benefit of safe choices capitalizing on past experience (exploitation) must be weighed against the potential benefit of risky choices flouting precedent (exploration) in order to extend or update this experience. Further tensions arise when a “cost of time” creates an urgency for reaching timely decisions [20–22]. For example, visual perception generates transiently stable interpretations of continuous sensory input and updates these interpretations at short intervals. Here, the benefit of remaining committed to an interpretation until analysis is complete must be weighed against the benefit of promptly overturning this interpretation in response to ongoing changes in the input. This tension reflects the competing objectives of perceptual stability and sensitivity [22, 23].

To study the inherent dynamics of sensory perception – as well as the trade-offs between conflicting objectives that this may entail – a suitable starting point is “multistable perception”, where perceived appearance perpetually changes at irregular intervals, shifting abruptly between distinct alternatives [24]. Presumably, this intrinsic instability reflects a spontaneous reassessment of perceptual decisions, even when the sensory scene remains unchanged [25, 26]. Numerous properties of multistable perception support this notion; notably, the ecological plausibility of alternative appearances and their dependence on behavioural context and prior experience [27]. Interestingly, the dynamics of multistable perception differs substantially between individuals of different age [28–33], and also in individuals with psychiatric disorders, such as autism spectrum disorder [34–38] among many others [39–45]).

To quantify the dynamics of multistable perception in cohorts of different age and psychiatric health, we tracked spontaneous reversals of binocular rivalry that were induced by presenting different images to each eye (**Fig. 1**). Importantly, we reliably and accurately inferred spontaneous reversals of subjective (covert) eye dominance from objective (overt) eye movements, thereby avoiding the confounds and biases of volitional reports [46]. Briefly, participants dichoptically viewed two horizontally moving gratings through a mirror stereoscope, while we recorded optokinetic nystagmus (OKN) in one eye. The slow phases (smooth pursuit) of the OKN pattern revealed which direction of movement was consciously perceived at the time [47–49].

**Figure 1:**
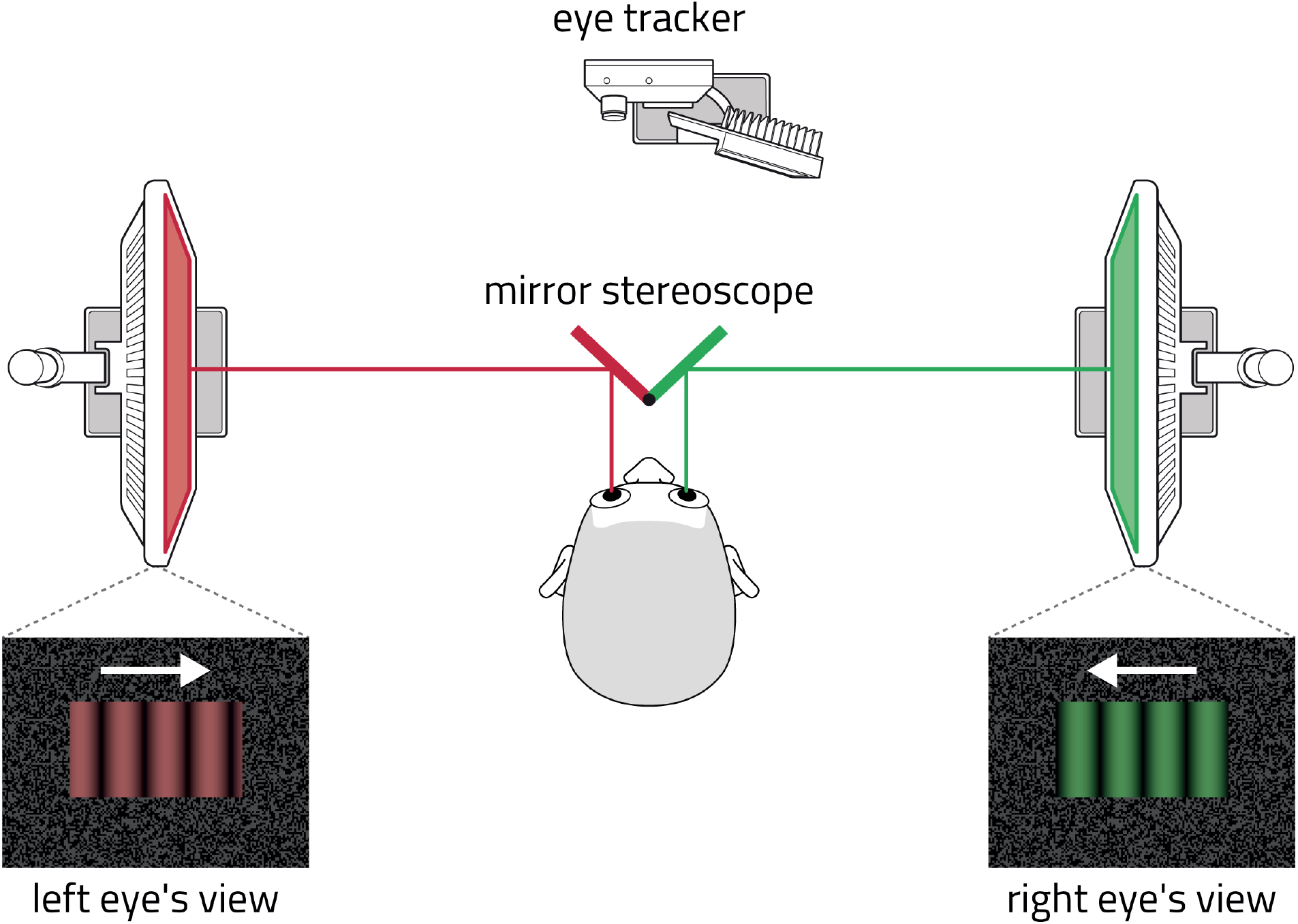
Binocular rivalry paradigm. Participants sat in a mirror stereoscope and viewed two separate displays with their left and right eyes. Displays showed gratings of different colour (red and green) and opposite motion, inducing occasional reversals of phenomenal appearance (binocular rivalry). Perceived appearance and its reversals were monitored objectively by recording optokinetic nystagmus of one eye (OKN), using an automated analysis [46].

With this approach, we quantified and characterized binocular rivalry dynamics in a large group of neurotypical participants (N=107) of different ages, from 12-year-old children to senior adults. This broad range of ages was intended to capture both developmental changes during adolescence and maturational changes through adulthood, into senior age. Additionally, we studied two psychiatric groups associated with developmental, genetic and environmental risk factors: Autism spectrum disorder (ASD; all males), which has a prevalence of about 1% in the population, and is diagnosed four times more often in males than in females [50], and where patients are known to present an atypical dynamics of multistable perception [34–38]; and borderline personality disorder (BPD; all females), which has an estimated prevalence between 1.6-5.9%, and is diagnosed predominantly in female (around three times more often than male) adolescents and adults [50, 51]. To our knowledge, multistable perception has not been studied in BPD so far.

After establishing three observable variables of binocular rivalry statistics, we were surprised to find that females and males follow quite different developmental trajectories, with particularly pronounced differences during adolescence and menopause. The maturational peak was observed at 19 years for females and 24 years for males, consistent with the earlier puberty onset time seen in girls compared to boys [52]. The binocular rivalry statistics of both psychiatric patient groups fell well outside these typical maturational trajectories.

From the observed rivalry statistics, we inferred the changing balance between conflicting objectives, by fitting a computational model of rivalry dynamics [53]. Modelling the rivalry dynamics of each age or patient group allowed us to conduct extensive simulations and to predict perceptual performance in volatile and unpredictable environments. This extrapolated perceptual performance could then be quantified in terms of the conflicting objectives of “stability”, “sensitivity”, and disposition for “exploration”.

Inferred trajectories of performance objectives again differed substantially between females and males. During adolescence, females seemed to gain on all three objectives, reaching a “sweet spot” of high stability, sensitivity, and exploration in their early twenties, where they remained until menopause. In contrast, males attained their highest levels of sensitivity and exploration in early adolescence, subsequently retreating slightly from this peak. Concomitantly, their perceptual stability increased until the mid-twenties and gradually declined thereafter. Psychiatric patient groups differed substantially from age-matched controls, raising the possibility that atypical development due to developmental, genetic, or environmental risk factors alters the functional optimization of perceptual decisions.

## 2 Results

### 2.1 Females and males show different developmental trajectories of binocular rivalry dominance statistics

During the experiment, visual stimulation was dichoptic and consisted of two horizontally moving gratings, differing in direction of motion and in colour contrast (red-and-black or green-and-black). This stimulation induced optokinetic nystagmus (OKN) in the direction the grating that currently dominates perception, so that (covert) reversals of subjective dominance (as well as transitional periods) could be inferred accurately from (overt) eye movements [46, 48, 54]. As this method does not require volitional reports from observers, it is well suited for developmental and patient groups. Determination of reversal timing is similarly precise for all developmental and patient groups (full-width confidence interval of 215 ± 54 ms). Participants viewed the dichoptic stimulus for ten trials of 95 seconds duration. The neurotypical participants included 21 developing children (12 twelve-year-olds, 19 sixteen-year-olds) and 52 adults (aged 18-69) (see Methods for details).

We established the statistics of perceptual dominance periods from the recorded eye movements in terms of three distribution moments, as these are highly diagnostic about the stochastic accumulation of sensory information that underlies perceptual decisions [26, 55, 56] **Fig. 2ab** shows the combined trajectories of median duration (M), interquartile range (IQR), and medcouple (MC, a robust measure of skewness) of dominance period distributions, separately for female and male observers, computed with a log-normal weighted sliding average. Trajectories of individual distribution moments are shown in **Supplementary Fig. 1**.

**Figure 2:**
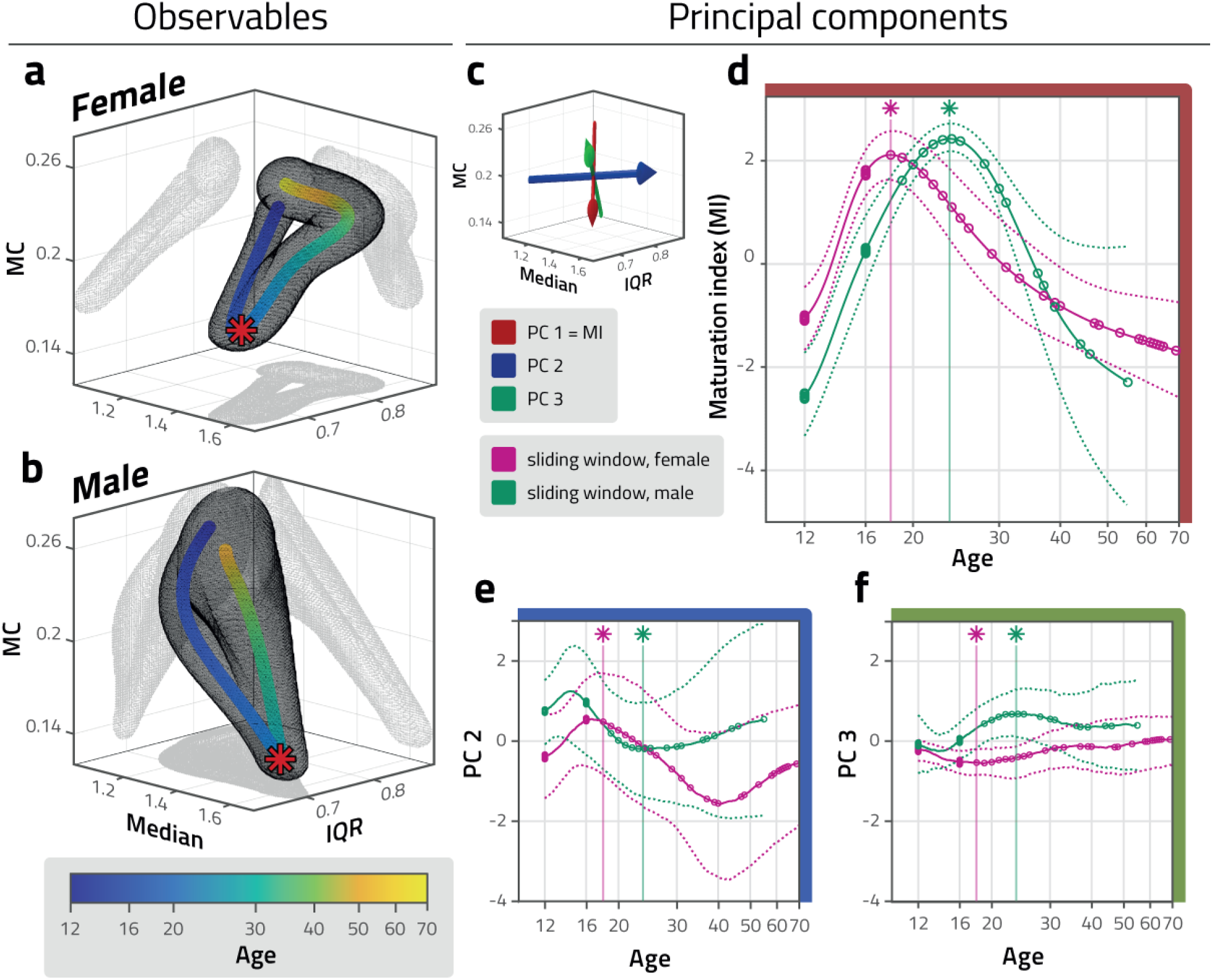
Developmental and maturational trajectory of distribution parameters. a, b: Parameters of distribution of dominance durations – median duration (median), interquartile range (IQR), and medcouple (MC) – during development and maturation of neurotypical female and male observers (blue to yellow colour scale). Red stars represent peak age of maturation index (d). The grey mesh represents mean values ± 50% SEM, computed with sliding log-normal weighting and repeated sampling with replacement. The developmental trajectories of the two sexes differ notably within this space. c: Principal component (PC) axes of mean distribution parameters observed over all ages and sexes (computed after conversion to z-score values). First (red), second (blue) and third (green) components account for 79%, 16%, and 3% of the variance, respectively. A maturational index (MI) is obtained by projecting mean parameter triplets onto the first principal component axis. d: Maturation index (MI) of neurotypical female and male observers (magenta and green, respectively), as a function of age. At age 18.9 and 23.7, peak values of 2.11 and 2.42 are reached by female and male observers, respectively (stars, vertical lines). Sliding window average (solid curves) and confidence intervals (±100% SEM, dotted curves) in harmonized units (z-score values). Individual observers are also indicated (circles). e, f: Development and maturation along second (e) and third (f) principal component axes, in harmonized units (z-score values). Note far smaller variance with age compared to (d).

As illustrated in **Fig. 2ab**, the combined trajectories of distribution parameter triplets M, IQR, and MC exhibit significant differences between females and males (age 12 to 70; *p* < 0.001). We assessed significance by comparing (differences between) random samples of female and of male observers with (differences between) random samples of ‘pseudo-female’ and ‘pseudo-male’ observers (see Methods for details). Even for partial trajectories, such as during early adolescence (age 12 to 14, *p* < 0.001), later adolescence (age 14 to 16, *p* < 0.001), young adulthood (age 14 to 24, *p* < 0.001), early maturity (age 24 to 40, *p* < 0.001), or later maturity (age 40 to 70, *p* < 0.01), the differences between females and males were significant. The respective starting points (age 12) of female and male trajectories differ significantly (*p* < 0.01), but the end points (age 50) do not (*p* ≈ 0.1).

When each distribution parameter is considered individually (Supplementary figure 1), lifetime trajectories of females and males are still significantly different (M with *p* < 0.01, IQR with *p* < 0.05, and MC with *p* < 0.01, assessed by comparing random samples of female and male observers with random samples of ‘pseudo-female’ and ‘pseudo-male’ observers).

Remarkably, the trajectory of every individual statistic (M, IQR, and MC) is U-shaped, in that the observed values reach an extremum in young adulthood (age 20), but in senior age (age 50 and over) return towards the levels observed in childhood (age 12) (see also **Supplementary Fig. 1**). The observed differences between children and young adults are statistically significant in all cases, for both sexes, and the same holds for differences between young adults and seniors (M with *p* < 0.01, IQR with *p* < 0.05, and MC with *p* < 0.01, based on random sampling with replacement).

This observation raises the question of whether lifetime trajectories are closed. In fact, the multivariate starting points (age 12) and end points (age 50) do not differ significantly for either female or male trajectories (*p* ≈ 0.2, based on random sampling with replacement). However, this may simply be due to the high degree of heterogeneity exhibited by senior observers. Thus, although starting and end points are clearly similar, we cannot conclude that they are identical.

### 2.2 Females peak earlier in maturation index derived from dominance statistics

To summarize the development of distribution parameters, we defined a maturation index (MI) by performing a principal component analysis on the z-scores of all distribution parameters (from both sexes). The first principal component accounted 79% of the variance. Thus, the projection of distribution parameter triplets onto this component, shown in **Fig. 2d**, summarizes the overall development and was used as a maturation index. The projection of parameter triplets on the second and third PCs accounted for 16% and 3% of the variance, respectively (**Fig. 2ef**).

In terms of index MI, females peak at an earlier age (age 18.9) than males (age 23.7). The difference in age is significant (*p* < 0.05) when random samples of female and male observers are compared with random samples of ‘pseudo-female’ and ‘pseudo-male’ observers (see Methods for details). The difference in peak height is not significant (*p* ≈ 0.1).

At the average age of ‘peak maturity’, the distribution parameter triplets M, IQR and MC exhibited by females and males differ significantly (*p* < 0.02). This is also true when female and male parameter triplets are compared at the individual peak value of the maturation index.

The lifetime trajectories of maturation index MI differ significantly between females and males (*p* < 0.01). This remains true also for partial trajectories, such as during adolescence (age 12 to 16, *p* < 0.01), young adulthood (age 18 to 24, *p* < 0.01),and early maturity (age 24 to 40, *p* < 0.001). No significant difference between females and males was obtained for some age cohorts that are represented less well in our sample (age 30 to 40, *p* ≈ 0.5; age 40 to 70: *p* ≈ 0.1).

Further differences in the lifetime trajectories of females and males are evident in terms of principal components PC2 (*p* < 0.05) and PC3 (*p* < 0.01). Of particular interest is principal component PC2, which changes the sign of its slope multiple times. At the female maturation peak (MI peak), the slope of PC2 changes from positive to negative (*p* < 0.01); whereas, at the male maturation peak, it changes from negative to positive (*p* < 0.01, based on random sampling with replacement). After the maturation peak, the slope changes once more around age 40 for females, being negative during ‘early maturity’ (age 26 to 40) and positive during ‘late maturity’ (age 40 to 50) (*p* < 0.01). For males, the slope after the maturation peak remains positive (*p* < 0.01).

Additionally, it seems possible that the slope of PC2 changes sign prior to the maturation peak (around age 14). However, this is not statistically significant (*p* ≈ 0.1), which may simply reflect the age structure of our male observer cohort.

### 2.3 Reinterpretation in terms of a computational model of reversal dynamics

To better interpret the results described above, we reproduced the particular dominance statistics of each group of participants with a computational model of binocular rivalry simulating a dynamic interaction of competition, adaptation, and noise [23, 53] (**Fig. 3ab**). Models of this type emulate important aspects of the cortical activity dynamics associated with binocular rivalry [57–60], namely, inhibitory interactions operating locally within visual representations [61–63], progressive weakening of the dominant representation by adaptation [64–66], and neural noise triggering reversals at irregular intervals [67–69]. However, the objective of such models is to reproduce lawful aspects of perceptual dynamics during binocular rivalry, not neurophysiological recordings. Besides dominance statistics (distribution moments) [23], models of this kind reproduce the stochastic resonance exhibited by binocular rivalry during harmonic counter-phase stimulation [68], as well as the counter-intuitive dependence of dominance durations on input strength (“Levelt’s propositions”) [26, 70, 71].

**Figure 3:**
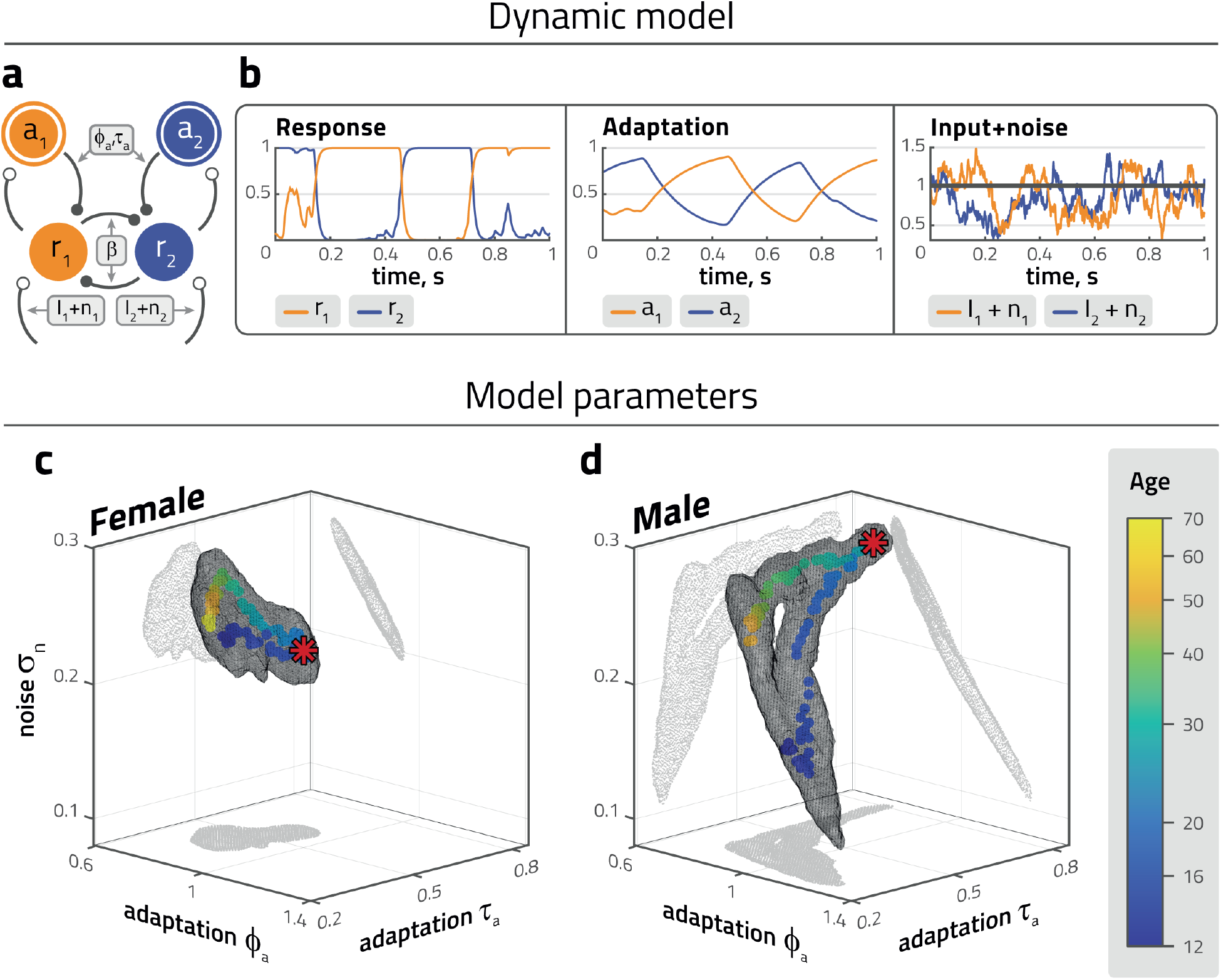
Development and maturation of fitted model parameters. a: Dynamic model of binocular rivalry, with competition, adaptation, and noise. Two representations (*r*_1_, *r*_2_) are driven by associated visual inputs (*I*_1_, *I*_2_) and independent noise (*n*_1_, *n*_2_). Each representation is inhibited by the other, as well as by the associated adaptive state (*a*_1_, *a*_2_). Free model parameters are competition strength (*β*), adaptation strength (*ϕ*_*a*_), adaptation time-scale (*τ*_*a*_), and noise amplitude (*σ*_*n*_). Input strength is fixed at *I*_1_ = *I*_2_ = 1. b: Representative example of model dynamics with abrupt dominance reversals of activities *r*_1_ and *r*_2_, gradual build-up and recovery of adaptive states *a*_1_ and *a*_2_ (middle), and noise *n*_1_ and *n*_2_ added to visual input *I* = 1 (*β* = 2, *ϕ*_*a*_ = 0.7, *τ*_*a*_ = 0.3, *σ*_*n*_ = 0.2). Reversals may be triggered by differential adaptive state *a*_1_ – *a*_2_, differential noise *n*_1_ – *n*_2_, or both. c, d: Parameter triplets *ϕ*_*a*_-*τ*_*a*_-*σ*_*n*_ fitted to reproduce (within a ±5%) experimentally observed mean distribution parameters. Competition strength was fixed at *β* = 3. Centroid values (coloured dots) and confidence range (± standard deviation, grey mesh). Red stars represent peak age of maturation index (**Fig. 2d**).

The four most consequential parameters of this model were strength of competition (*β*), adaptation strength (*ϕ*_*a*_), noise (*σ*_*n*_) (all dimensionless) and the time-constant of adaptation (*τ*_*a*_). Simulated rivalry reversals were generated with four levels of competition strength (*β* = 1, *β* = 2, *β* = 3, and *β* = 4). All values of *β* produced qualitatively similar results and illustrations show the results for *β* = 3. Input strength was fixed at *I*_1,2_ = 1. Other, less consequential parameters were fixed as well (time-constants of activity *τ*_*n*_ and noise *τ*_*r*_, inflection point of activation function *k*; see Methods). We systematically varied *ϕ*_*a*_, *τ*_*a*_, and *σ*_*n*_ in order to determine the volume in the *ϕ*_*a*_-*τ*_*a*_-*σ*_*n*_ subspace in which the average distribution moments exhibited by a particular age cohort of observers were reproduced within a tolerance of ±5% tolerance (see Methods for details).

The resulting volumes of model parameters in *ϕ*_*a*_-*τ*_*a*_-*σ*_*n*_ subspace are shown in **Fig. 3cd** for neurotypical female and male participants. Here, the previously observed average trajectories in terms of distribution moments (**Fig. 2ab**) are transformed into new trajectories in terms of model parameters. Although model fitting introduces a degree of stochasticity, the transformed trajectories exhibit significant differences between females and males over the entire lifespan, e.g., during early adolescence (age 12 to 14, *p* < 0.001), middle adolescence (age 14 to 16, *p* < 0.01), late adolescence (age 16 to 18, *p* < 0.001), at the respective maturational peaks (females aged 18 to 20, males aged 22 to 25, *p* < 0.001), early maturity (age 30 to 40, *p* < 0.001), middle age (age 40 to 50, *p* < 0.001) and later maturity (age 50 to 60, *p* < 0.001).

Females and males develop similarly with regard to adaptation time constant *τ*_*a*_. In both sexes, *τ*_*a*_ increases from childhood until the maturational peak, only to decrease during subsequent maturation. Distinctly, females develop toward a maturational peak with slightly larger values of adaptation strength *ϕ*_*a*_, and slightly smaller values of noise *σ*_*a*_, while males develop towards a maturational peak with considerably smaller values of adaptation strength *ϕ*_*a*_, and considerably larger values of noise *σ*_*a*_. The development of females and males after their maturational peaks is also distinct: at age 40, females develop larger levels of noise and smaller levels of adaptation than males.

It is unclear whether male and female parameter trajectories are fully closed. As mentioned above, the distribution moments observed in childhood and in old age do not differ significantly (either for females or for males). Additionally, a trade-off between adaptation and noise introduces an ambiguity in model fitting, which increases toward small values of *τ*_*a*_ (i.e., the confidence range represented by the gray mesh is rather large).

### 2.4 Trajectories of perceptual objectives predicted for volatile environments

A second goal of this study was to characterize the perceptual performance of female and male observers of different ages in volatile and unpredictable sensory situations, in order to understand how these groups balance conflicting perceptual objectives such as stability, sensitivity or exploration. Unfortunately, it is not feasible to empirically establish a correlation between stochastic sensory stimuli and stochastic perceptual choice responses, as there would be far too many joint possibilities to consider. However, given a dynamic model for the perceptual choice response, responses to stochastic sensory stimuli can be predicted from exhaustive simulations (approximately 30 hours of simulated viewing time). Of course, the value of such predictions hinges on the validity of the model.

To predict perceptual performance of observers in a volatile environment, we simulated approximately 106 perceptual reversals for any given dynamic model (i.e., any particular set of model parameters), in response to stochastically time-varying inputs *I*_1_ and *I*_2_. Specifically, while mean input (*I*_1_ + *I*_2_)/2 remained constant, input bias Δ*I* = *I*_1_ − *I*_2_ varied as a continuously stochastic process (see Methods for details). This allowed us to assess the relative influence on perceptual reversals of the sensory environment (external state *I*_1_, *I*_2_), on the one hand, and of the internal perceptual dynamic (internal state variables *r*_1_, *r*_2_ and *a*_1_, *a*_2_), on the other hand. At any given moment, the model responded to these external and internal factors either with ‘perceived stability’ (continued dominance) or ‘perceived novelty’ (reversal of dominance).

Further details of these *in silico* experiments are illustrated in **Fig. 4a**. Sensory input bias Δ*I* = *I*_1_ – *I*_2_ varied as a continuously stochastic process with constant mean, variance, and autocorrelation (see Methods). The superposition of sensory input *I*_1_, *I*_2_ and intrinsic noise *n*_1_, *n*_2_ are also shown. The internal state variables of levels of activity (*r*_1_, *r*_2_) and levels of adaptation (*a*_1_, *a*_2_) respond dynamically and stochastically. Activity levels *r*_1_ and *r*_2_ assume mostly categorical values (near zero or near unity), with intermediate values occurring only transiently during reversals (defined as crossover points with *r*_1_ = *r*_2_). In contrast, adaptation levels *a*_1_ and *a*_2_ change gradually over time and their combined state may be summarized conveniently in terms of adaptation bias Δ*a* = *a*_1_ − *a*_2_. Perceptual performance in this stochastic environment is encapsulated in the joint statistics of input bias Δ*I*, adaptation bias Δ*a*, and model response (reversal or no reversal).

**Figure 4:**
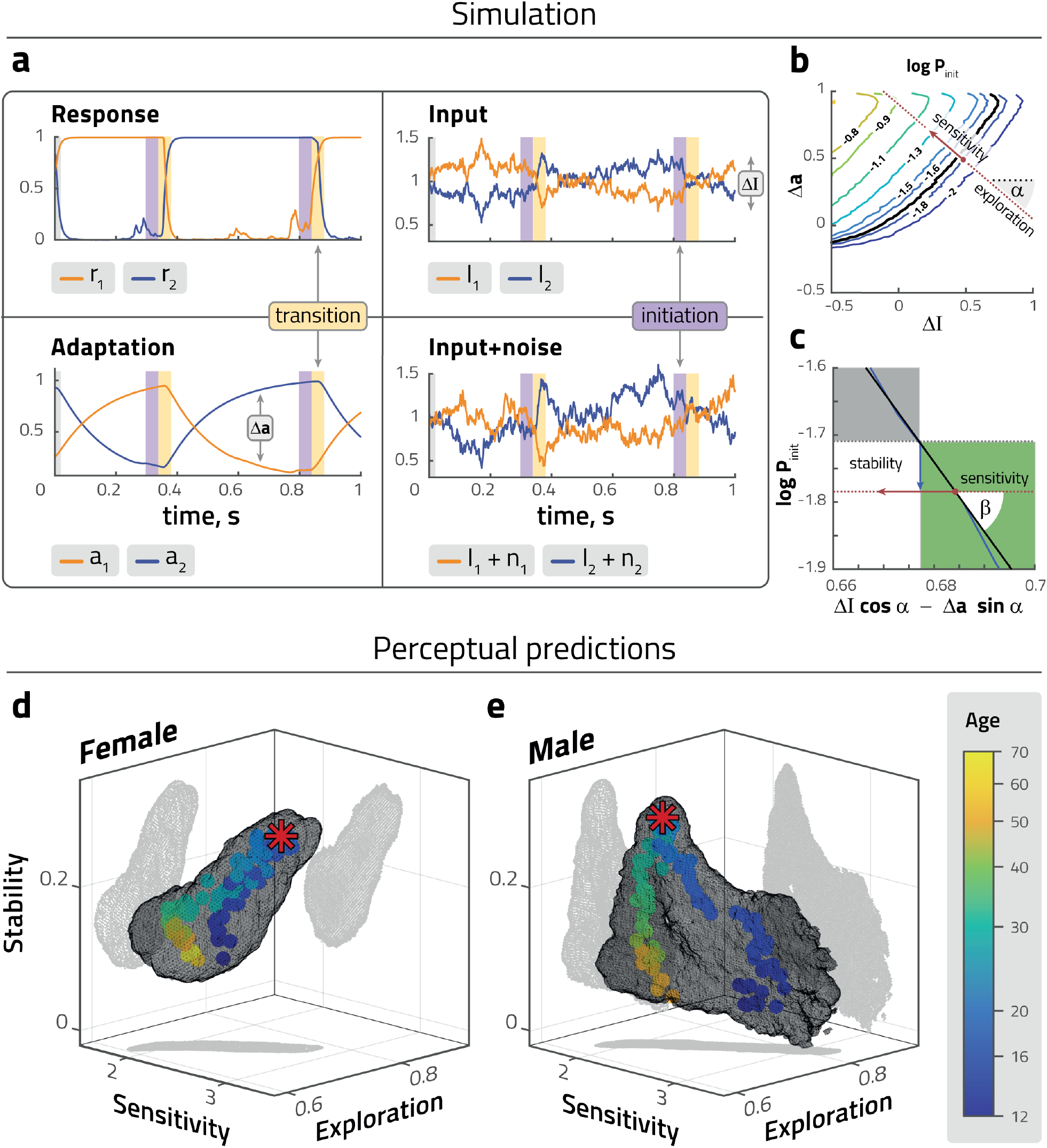
Predicted perceptual behaviour in a volatile environment. a: Representative example of simulated dynamics of fitted model: activities *r*_1_, *r*_2_ and adaptive states *a*_1_, *a*_2_ driven by intrinsic noise *n*_1_, *n*_2_ and variable sensory inputs *I*_1_ and *I*_2_. In a volatile environment, reversals (defined by *r*_1_ = *r*_2_) may be triggered externally (differential input Δ*I* = *I*_1_ *− I*_2_) or internally (differential adaptive state Δ*a* = *a*_1_ − *a*_2_) or both. Competition strength was fixed at *β* = 3. To assess reversal initiation, we distinguished between transition periods (yellow stripes, 20ms before and after *r*_1_ = *r*_2_), immediately preceding periods (purple stripes, 40 to 21 ms before *r*_1_ = *r*_2_), and all other times. b: Based on this classification, we computed the conditional likelihood of reversals (coloured contours) as a function of Δ*I* and Δ*a* values during initiation periods, in the vicinity of the median Δ*I* and Δ*a* values (over all periods, red dot). In this vicinity, the reversal probability grows exponentially in a particular direction (red dashed line). The length of the gradient vector *∂* ln *P* (Δ*I*, Δ*a*) represents the ‘sensitivity’ of reversal probability to Δ*I* and/or Δ*a* values, and the ‘exploration’ angle *α* represents the relative influence of internal state Δ*a*, compared to external state Δ*I* (external state). A larger value implies that the system responds less consistently to external state, behaving in a more explorative manner. c: Enlarged cut through the ln *P* (Δ*I*, Δ*a*) surface in the direction of the the gradient vector (red arrow), showing the neutral level (defined by Δ*I* = Δ*a* = 0, black dashed line) and median level (red dashed line). The distance between levels (blue arrow), represents ‘stability’ and measures the stabilizing or destabilizing effect of external and internal median states. For positive values (within green area), median states lower reversal probability, stabilizing the percept; vice versa for negative values (within grey area). d, e,: Developmental and maturational trajectories of predicted perceptual parameters (stability, sensitivity, exploration) for females (d) and males (e). Mean values (coloured dots), peak age of maturation index (red stars), and confidence intervals (standard deviations, grey volumes).

In order to focus the analysis on the period of initiation or non-initiation of reversals, we disregarded the largely stereotypical time course of reversals themselves. Specifically, we excluded transition periods (20 ms before and after each crossover point) and analyzed both initiation periods (defined as 40 ms to 21 ms before a crossover) and non-initiation periods (all other times) at sampling intervals of 1 ms. The analysis provided a large ensemble of value pairs of Δ*I* and Δ*a* from initiation periods and an even larger ensemble of value pairs from non-initiation periods. To characterize the stochastic dependence of reversal initiation on external state Δ*I* and internal state Δ*a*, we computed the conditional probability *P*_*init*_ (Δ*I*, Δ*a*) of reversals as a function of Δ*I* and Δ*a*. The accuracy of this probability surface was highest in the vicinity of the median state Δ*I*_*m*_, Δ*a*_*m*_, where sampling density was largest (**Fig. 4b**; see Methods for details).

The results showed that the logarithm of initiation probability ln *P*_*init*_ varied almost linearly with state variables Δ*I* and Δ*a* (linear regression *r*^2^ > 0.99) in the vicinity of the median state (Δ*I*_*m*_, Δ*a*_*m*_). Note that ln *P*_*init*_ grows with negative Δ*I* and positive Δ*a* (because positive input bias favours the dominant, and positive adaptation bias the suppressed percept). As a planar surface, the function ln *P*_*init*_(Δ*I*, Δ*a*) had three degrees of freedom, which corresponded to three different perceptual objectives: *sensitivity*, *exploration*, and *stability* (**Fig. 4bc**). The motivation for this particular choice of terms is provided in the Discussion and in Methods. Specifically, *sensitivity* represents the maximal gradient of ln *P*_*init*_ with respect to state variables Δ*I* and Δ*a*. Mathematically, it was defined as the *length* of the gradient vector *∂* ln *P*_*init*_(Δ*I*, Δ*a*) at the median state (or, equivalently, as slope angle *β*). Similarly, *exploration* represented the relative influence of internal state Δ*a*, compared to external state Δ*I*, and was defined mathematically as the *direction* of the gradient vector *∂* ln *P*_*init*_(Δ*I*, Δ*a*) at the median state (or, equivalently, as angle *α*). Finally, *stability* measures the stabilizing effect of reversals. Mathematically, it was defined as the difference between ln *P*_*init*_ at the median state (Δ*I*_*m*_, Δ*a*_*m*_) and the neutral state (Δ*I* = 0, Δ*a* = 0), where any reversals are due to external or internal noise. Typically, ln *P*_*init*_ is smaller in the median state than in the neutral state, because reversals bring about a more stable situation (**Fig. 4c**).

In this way, every set of model parameters *β*-*ϕ*_*a*_-*τ*_*a*_-*σ*_*n*_ that reproduced the reversal statistics of a particular group of participants could be mapped onto a corresponding value triplet stability-sensitivity-exploration of perceptual objectives. The resulting trajectories of stability, sensitivity, and exploration associated with different age groups are shown in **Fig. 4de**. Although this procedure introduced additional stochasticity, notable differences between the developmental and maturational trajectories of the sexes are evident over the entire lifespan, i.e. during early adolescence (age 12 to 14, *p* < 0.01), middle adolescence (age 14 to 16, *p* < 0.01), late adolescence (age 16 to 18, *p* < 0.001), at the respective maturational peaks (females aged 18 to 20, males aged 22 to 25, *p* < 0.001), early maturity (age 30 to 40, *p* < 0.05), middle age (age 40 to 50, *p* < 0.05) and later maturity (age 50 to 60, *p* < 0.05).

Females gained higher values of stability, sensitivity, and exploration during development, reaching maximal values of each parameter at the maturation peak, with values subsequently decreasing during maturation. Males, on the other hand, exhibit their peak values of sensitivity and exploration during adolescence (around age 14). As they approach the maturational peak, they increase stability while decreasing sensitivity and exploration. During subsequent maturation, stability decreases while sensitivity and exploration remain approximately stable. These results hint at different developmental strategies between sexes, in that females gained steadily on all three objectives – stability, sensitivity, and exploration – during adolescence until young adulthood (maturational peak), while adolescent males start out with high sensitivity and exploration, only to subsequently gain stability at the expense of sensitivity and exploration until young adulthood (maturational peak).

### 2.5 Clinical populations fall outside of typical developmental and maturational trajectories

In addition to neurotypical observers of various ages, we performed the same set of procedures - binocular rivalry experiment, model fitting, and volatile-environment simulations - with two adult clinical groups: 12 females with borderline personality disorder (BPD; mean age 27.1), and 12 males with autism spectrum disorder (ASD; mean age 28.5). The observed binocular rivalry dominance distribution parameters, and predicted perceptual parameters are shown in **Fig. 5**, contrasted with the corresponding sex’s typical developmental and maturational trajectories (results of the model fitting are shown in **Supplementary Fig. 1**).

**Figure 5:**
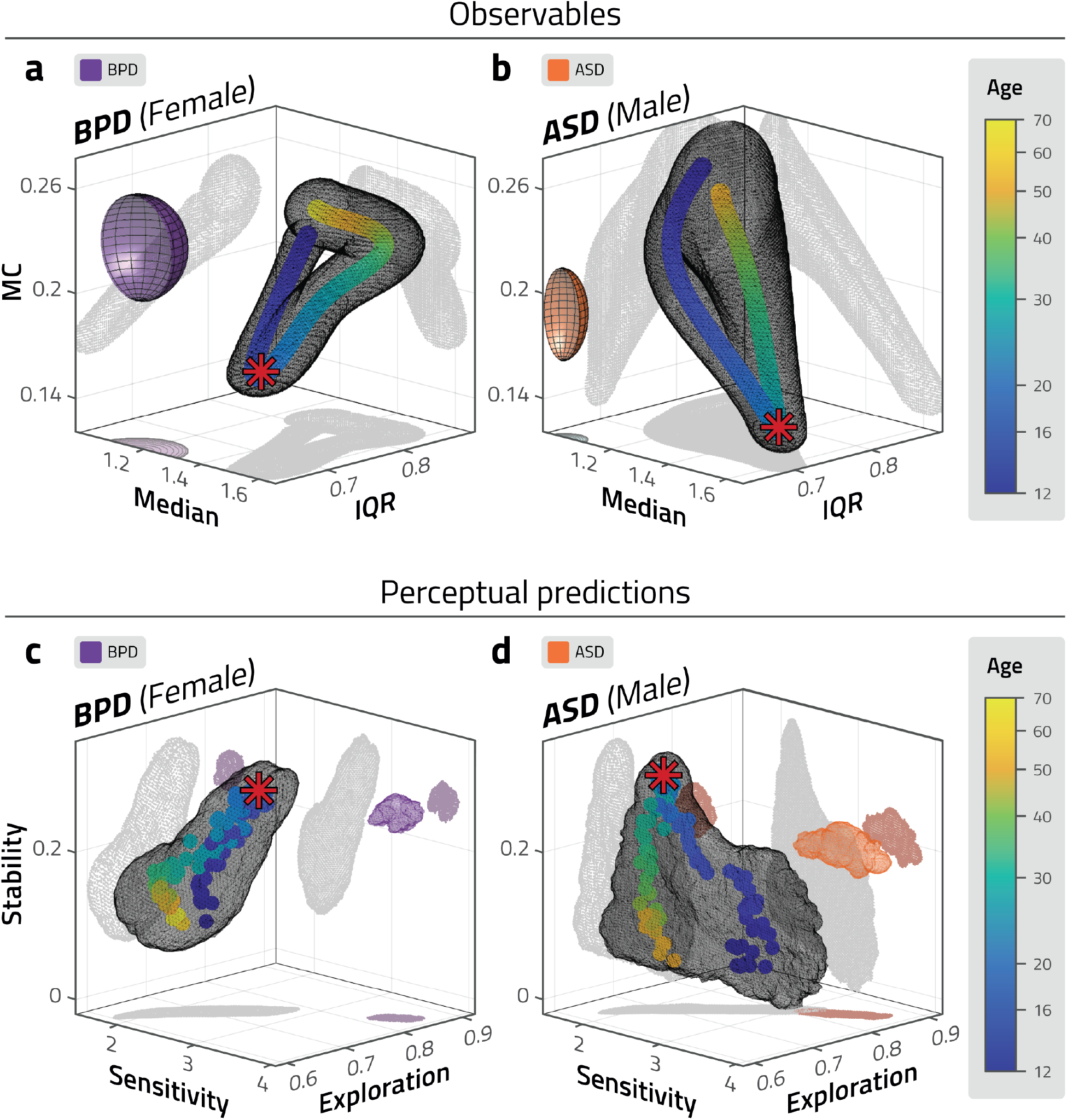
Observed distribution parameters and predicted perceptual behaviour of atypical groups. Observed distribution parameters (a, b), and predicted perceptual behaviour in volatile environments (c, d) for atypical groups, compared to neurotypical trajectories. Observers with borderline personality disorder (BPD, all female) are compared to neurotypical females of different ages. Observers with autism spectrum disorder (ASD, all male) are compared to neurotypical males of different ages. (a, b) Ellipsoids represent mean SEM values for BPD (purple) and for ASD (in orange). (c, d) Coloured volumes represent confidence intervals (standard deviation) for BPD (purple) and for ASD (in orange). The dominance period statistics and perceptual predictions of both groups fall outside the typical developmental and maturational trajectory.

The results show that both atypical groups fall outside the typical developmental and maturational trajectories. Despite the different character of their respective disorders, they differ from the typical trajectories in similar ways. In terms of observable dominance distribution parameters (**Fig. 5ab**), both groups show lower median and IQR, and higher MC values. In the predicted perceptual performance in a volatile environment (**Fig. 5cd**), both groups have higher values of sensitivity and exploration, compared to neurotypicals.

## 3 Discussion

The main purpose of our study was to establish lifespan trajectories of visual perceptual behaviour. With respect to neurotypical brain development and maturation, our results indicate that visual decision making, at least insofar as this is reflected in multistable perception, changes markedly in adolescence, and then more gradually across the lifespan. The fact that the lifetime trajectories we found were sex-specific, suggests that biological maturation plays a major role in visual decision making. Neuroendocrine changes, especially levels of sex steroid hormones, are known to pace bodily growth, emotional development, and cortical pruning in adolescence [5, 72, 73]. Neuroendocrine changes also seem to influence cognitive function during [74, 75] and after the teenage years [76, 77]. The fact that the maturation index in our study peaks earlier in females (around 19 years of age) than in males (around 24 years of age) is consistent with the relative onset times of puberty in girls and boys [52]. The comparative lateness of these maturational peaks may also be significant. For example, it may be that visual decision making is a neotenous function in humans, which involves the latest maturing brain areas, such as prefrontal cortex and precuneus [78, 79]. Looking further into maturation, after the apex of young adulthood, males exhibit a progressive decline of sex hormones [80], whereas females experience a sharp gonadal hormone falloff later in life [81]. Interestingly, we observed a significant change in the direction of the maturational trajectory of females around age 40. No such directional change was evident in males.

The observed statistics of binocular rivalry periods – median duration (M), interquartile range (IQR), and medcouple (MC) – reveal the dynamic accumulation of sensory information that underlies visual decision making [26, 55]. However, by themselves, these statistics are not easily interpreted and any implications for visual decision making are not immediately evident. To generate an interpretive hypothesis, we sought to predict the perceptual behaviour of observers in volatile and unpredictable sensory situations by means of extensive simulations (approximately 30 hours of simulated viewing time). Such extensive data would have been difficult to obtain with human observers.

As a first step, we fitted a stochastic dynamic model to reproduce the reversal statistics (i.e., M, IQR and MC of rivalry periods) of a particular group of human observers (e.g., females aged 25 years). In a nutshell, this model categorizes external input by choosing between two internal stable states. If the input changes, an ‘external’ tension (mismatch) between internal and external states may develop. Additionally, the model actively builds ‘internal’ tension in terms of an adaptive state which contradicts its internal state. Occasionally, the model relieves tension by executing a reversal, bringing external, internal, and adaptive states back into closer correspondence.

As a second step, we characterized the perceptual behaviour of this model when image contrast varied unpredictably over time. This simulated behaviour was then summarized quantitatively by three measures. The first measure, *stability*, captured the stabilizing effect of reversals. It reflects the degree of stability that the dynamic system (by virtue of its stable states) imposes on volatile external inputs. The second measure, *sensitivity*, reflects how much reversal probability changes with rising tension between internal and external states. The third measure, *exploration*, describes the relative influence of internal tension (differential adaptation) compared to external tension (differential input).

Although defined in the context of a stochastic dynamic model, these three measures are broadly analogous to competing objectives of perception prescribed by the framework of reinforcement learning. When objects can appear or disappear at any time, the thoroughness of object classifications must be balanced against the frequency of such classifications [20, 21]. In our context, this corresponds to the tension between *stability* (temporal persistence of current choice) and *sensitivity* (susceptibility to changed circumstances, internal or external) [22, 23]. Additionally, in an unstable world, the value of past experience diminishes with time, so that the benefit of past experience must be weighed against the benefit of discovering something new (exploitation-exploration dilemma [17–19]). In our context, this corresponds to the influence of external states (input bias), relative to internal states (adaptation bias) that prompt a fundamental reassessment of such evidence. Hence the relative weight of adaptation bias represents a tendency for *exploration*.

In terms of behavioural measures predicted by our model, females are characterised by increasing values of stability, sensitivity, and exploration during adolescent development. Peak values of all three measures are reached by twenty, defining a sweet-spot of optimal function for perceptual decisions. After the peak, a moderate decline can be observed until menopause, which is similar in all three parameters. Signs of a stability-sensitivity trade-off can be observed after menopause when stability declines sharply, with slight increases in sensitivity and exploration, following the male pattern in this age-group. Males, on the other hand, have peak values of sensitivity and exploration by the age of sixteen, exceeding the highest female levels. The heightened sensitivity is coupled with low levels of stability, demonstrating a clear trade-off between these measures, at least at the extreme end of the range. Until the mid-twenties, stability rises at the expense of sensitivity and exploration. The male sweet-spot of optimal function seems to be characterised by lower sensitivity and exploration levels, but a higher level of stability, when compared to females. After the peak, males decline progressively in stability. This intriguing and complex pattern in the lifespan evolution of the two sexes suggests, on one hand, that the rules of growth are not uniform even in the developmental period, and on the other hand, that while parameter values in senior age are similar to those in childhood, ageing is not simply the “reverse” of development, as it has been suggested with respect to other cognitive functions as well [82].

Given the biological determination reflected in the age- and sex-specific differences, we interpret this pattern of findings as a result of alternative developmental strategies. It seems that females approach the peak in all three parameters faster, and although this provides them with lower peaks as compared to males, it also seems to provide for a greater stability throughout the childbearing years, with a major decline only after menopause. The trade-off between stability and sensitivity is more obvious in males who start with maximum levels of sensitivity and exploration mid-adolescence, and in parallel to stability building up by the mid-twenties, a great deal of sensitivity is lost. The greater exploration range in the early years, the higher apex, and the continuous decline after the apex might indicate a developmental strategy to fine-tune the individual for the age of highest fertility in males [83]. These alternative developmental strategies suggested by our simulated experiments - one focusing on stability throughout the childbearing period, and the other, focusing on highest performance by the peak fertility age - are particularly interesting in the context of perceptual decisions. Decisions made between alternatives, even if not volitional or highly conscious, make up our everyday life, and the particular style with which we are dealing with those decisions may have crucial effects on our lives. It makes a difference, for example, whether our brain is tuned to be relatively insensitive to environmental changes, rendering decisions stable and stereotypical and preventing extensive exploration or, in another scenario, whether higher sensitivity is combined with a heightened inner drive for exploration.

After obtaining developmental and maturational lifetime trajectories of neurotypical subjects, it is of particular interest how these would be drawn when biological conditions or environmental factors are not adequate for typical development. To this end, we have tested two psychiatric groups: adult participants with autism spectrum disorder (ASD, all males) and with borderline personality disorder (BPD, all females). Both the experimentally observed distribution parameters and the predicted behavioural measures fell completely outside of the typical developmental trajectories for their respective sex –– i.e., adults with these disorders differed not only from neurotypical adults of a similar age, but from the measures of any examined ages, probably missing the sweet-spot of optimal function at an earlier age. In terms of the trade-off in predicted behavioural measures, both patient groups demonstrate sensitivity and exploration levels beyond the range of neurotypicals, while stability is reduced markedly only in males, resembling the pattern seen in neurotypicals. As all psychiatric patients in the study were adults, we cannot pinpoint the when-and-how of the deviations from typical trajectories in ASD and BPD from these results. It is likely that the onset of deviations is present from early childhood in ASD as sensory symptoms and differences in visual perception are already present in childhood [84], and multistable perception already differs from that of typically developing children before adolescence [37]. Since BPD is not characterised as a neurodevelopmental disorder, deviations probably have a later onset. By adolescence, the disorder can be reliably diagnosed [51], so the deviations from typical trajectories likely occur during, or somewhat before adolescence.

In interpreting the clinical findings, it should be mentioned that BPD is a disorder that predominantly affects females [50] (although one study found equal prevalence of BPD among both sexes [85]). Distinctive patterns in hormone levels, especially the relative changes in ovarian hormones may induce the expression of BPD features [86]. Our BPD participants seem to be in a higher sensitivity and exploration range than neurotypical women of the same age, however, this is obtained at the expense of stability, demonstrating the force of the trade-off, especially outside of the sweet-spot. With respect to ASD which is more often diagnosed in males [50], a similar tendency can be observed: increased levels of sensitivity and exploration and reduced stability as compared to neurotypicals of similar age. Although female-underdiagnosis [87] should not be overlooked, a popular theory of autism claims that the autistic brain is a hypermasculinized version of the male brain due to increased fetal testosterone levels [88]. In terms of excessive levels of sensitivity and exploration, our result from simulated experiments support this picture, although alternative interpretations, such as a general drawback of sex differentiation [89, 90] cannot be ruled out. Including male BPD, female ASD participants, larger samples of patient groups, and hormonal assessments would be essential in further studies.

Our results regarding ASD and BPD may inform the field of computational psychiatry, which aims at the large-scale phenotyping of human behaviour using computational models, with the hope that it may structure the search for genetic and neural contributions of healthy and diseased cognition [91]. Within this field, the framework of developmental computational psychiatry aims to establish normative developmental trajectories of computations, relate them to brain maturation, and determine when and how they deviate in mental disorders, in order to help uncover the relationship between the changes of brain organization in childhood and adolescence, and the heightened vulnerability to psychiatric disorders in these periods [15]. Our results fit into this framework by establishing typical developmental trajectories of visual decision making, and relating results from subjects with mental disorders to these trajectories. They also may serve as a small step of the large-scale phenotyping efforts to better understand the nature of mental disorders in terms of aberrant computations.

To conclude, in our detailed study on the lifespan trajectories of perceptual performance employing a no-response binocular rivalry paradigm combined with dynamic computational modelling, we have found characteristic age- and sex-specific developmental and maturational trajectories, with marked differences between neurotypical and psychiatric populations. These trajectories should serve to better describe our own neurocognitive phenotype and reveal relevant factors behind atypical development underlying mental health disorders.

## 4 Methods

### 4.1 Participants

A total of 107 participants took part in the binocular rivalry experiment: 28 twelve-year-old (19 female), and 19 sixteen-year-old (10 female) children; 52 neurotypical adults (average age 35.9, range 18 to 69, 32 female); 12 adults with autism spectrum disorder (ASD, average age 29, range 19 to 44, n=12, all male), and 12 adults with borderline personality disorder (BPD, average age 27, range 20-37, n=12, all female). Participants were considered typically developing, or neurotypical, if they reported no history of mental illness or disorder.

Nine of the twelve participants with ASD were recruited from the Department of Psychiatry and Psychotherapy, Semmelweis University. These participants were diagnosed by a trained psychiatrist. They went through a general psychiatric examination, and their parents were interviewed about early autism-specific developmental parameters. All participants fulfilled the diagnostic criteria of ASD, including autism-specific signs between the ages of 4-5 years. The other three participants in this group were recruited from Aura Organization, a nonprofit organization assisting people with ASD. We did not collect further diagnostic information from these participants. The participants with BPD were all recruited from the Department of Psychiatry and Psychotherapy, Semmelweis University. Their diagnostic status was assessed by the Hungarian version of the Structured Clinical Interview for the Diagnostic and Statistical Manual of Mental Disorders, fourth edition, Axis I and II disorders.

All participants had normal or corrected to normal vision, and reported no colour blindness. Before the experiment, all participants passed a stereoacuity test (Super Stereoacuity Timed Tester, by Stereo Optical Co., U.S. Patent No. 5,235,361, 1993). All adult participants, and all the children’s caregivers have provided informed written consent, as well as the parents or legal guardians of the subjects with ASD, where applicable (i.e. where subjects were under guardianship). The experiment was conducted in accordance with the relevant guidelines and regulations for research involving human subjects, and was approved by the Ethical Review Committee of the Institute of Psychology, Pázmány Péter Catholic University for neurotypical participants, and by the Semmelweis University Regional and Institutional Committee of Science and Research Ethics for participants with a psychiatric condition. Participants were given a book voucher for their participation.

### 4.2 Binocular rivalry experiment

Participants were fixated at a headrest during the experiments (SR Research Head Support, https://www.sr-research.com). The setup for dichoptic stimulation consisted of two LCD displays (subtending 26.6° horizontally and 21.5° vertically, with an approximate resolution of 48 pixels/° of visual angle, and a refresh rate of 120 Hz), which participants viewed through two 45° mirrors, attached to the headrest. The mirrors were coated, such that they reflected the visible light spectrum, but transmitted infrared light, allowing the use of an infrared camera for optical eye tracking.

The participants viewed green-and-black gratings with one eye, and red-and-black gratings with the other eye. The gratings moved horizontally, either consistently (in the same direction), facilitating perceptual fusion, or inconsistently (in opposing directions), facilitating perceptual rivalry. Each grating subtended a rectangular area of 15.2° width and 8.4° height. The spatial frequency was 0.26 cycles/°, and the temporal frequency 8.7 cycles/s. The motion’s speed was 33.5°/s or 1600 pix/s. Gratings were framed in a rectangular box with a random texture pattern, in order to facilitate binocular fusion. Stimuli were generated with Psychophysics Toolbox 3 [92–94] running under MATLAB R2015a. The display’s spatial resolution was 48 pix/°, and its temporal refresh rate was 120 Hz.

Before the experiment, participants were asked to view the display as attentively as possible, and to follow the horizontally moving gratings with their gaze. This was introduced with analogies such as “follow them like you would follow passing trees on a moving train”. We did not tell them that they will see rivalling stimuli, only that if the direction of the gratings they see will seem to change, let their gaze change direction too. We asked them to refrain from blinking as much as convenient. Participants did not have to report which stimuli they were perceiving at any time. Instead, perceptual states and transitions were calculated from eye-movement recordings.

The experiment consisted of ten trials, each 95 s long. The initial trial (introductory trial) served to familiarize participants with the display, and was not included in the analysis. It started with 22 s of consistent grating motion in alternating directions, followed by 72 s of inconsistent motion, and finishing with 1 s of consistent motion. During the introductory trial, we provided feedback for the participants on the behaviour of their eye movements. The following nine trials (experimental trials) began with 2 s of consistent motion, followed by 92 s of inconsistent motion, and ended with 1 s of consistent motion. The consistent episodes served to reduce eye strain and to test the ocular response to physical motion reversals. On experimental trials, participants received no feedback on their behaviour. Across trials, the colour (either red or green) and direction (either leftward or rightward) shown to each eye was altered. The experiment consisted of three blocks. After the third and sixth experimental trials, participants had a 5-minute break.

### 4.3 Establishing reversal sequences and dominance statistics from OKN

When a rivalrous display induces horizontal OKN, the direction of smooth pursuit phases provides a reflex-like indication of perceived direction [47–49]. During the experiment, we recorded horizontal eye position of subjects with a sample rate of 1000 Hz, and inferred reversals of perceived direction with the cumulative smooth pursuit (CSP) method, described elsewhere [46]. Briefly, the method removes off-scale values (blinks and other artefacts), and defines slow pursuit segments by a compound criterion (slow velocity |*v*| ≤ 1.5 pix/ms, low acceleration |*a*| ≤ 0.12 pix/ms^2^, duration > 50 ms), aligns slow pursuit segments, interpolates gaps, then subsamples and fits the resulting sequence multiple times ( “bagging” [95]). The result of this robust splining procedure is estimated eye velocity (median and 95% CI) at every time-step. Dominance periods were defined as contiguous intervals in which the entire 95% CI is either above or below a gaze velocity threshold of ±0.1 pix*/*ms. All other intervals were designated as perceptual transition periods.

As the rate of perceptual reversals often accelerates while viewing a binocular rivalry display [96, 97], the initial 30 seconds of each trial were discarded from analysis. We pooled the remaining reversal sequences obtained for each observer across different trials, and calculated the median (M), interquartile range (IQR), and medcouple (MC) of dominance durations. These robust statistical measures were used to reduce the effect of outliers. Median and interquartile range (IQR) provide robust alternatives for first and second moments (mean and variance), while medcouple (MC) offers a robust alternative for the third moment (skewness), which is particularly sensitive to outliers.

In previous work [46], we reported the latency of reversal detection to be approximately 180 ms, with a 95% confidence interval of 97 ms. This compares favourably to volitional reports, which showed an average latency of approximately 450 ms, with a confidence interval 150 ms.

Although oculomotor parameters do change with age, this did not bias our results. The full-width 95% confidence intervals for reversal timing was approximately 215 ms for a ll observer cohorts (developmental and patient), with a standard deviation of approximately 54 ms. Specifically, confidence intervals for different cohorts were 205 ± 51ms (age 12), 215 ± 57ms (age 16), 223 ± 55ms (age 25), 205 ± 51ms (over 50), 224 ± 56ms (BPD), and 223 ± 57ms (ASD).

### 4.4 Establishing developmental and maturational trajectories

For each observer *i*, three distribution parameters were established as described above: *M*_*i*_, *IQR*_*i*_, *MC*_*i*_, and age *a*_*i*_. From the individual observer values, average values were computed separately for female and male observers. As development slows down with age, sliding averages were computed with log-normal weighting, so that window size increased proportionally with age. For median 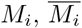 was computed as follows:

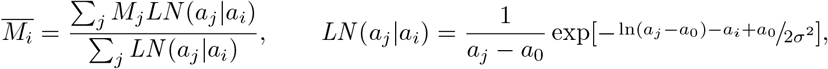

where *a*_0_ = 8 and *σ* = 0.35. 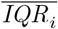 and 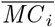 were obtained analogously.

Confidence intervals were computed by repeatedly sampling observers (of a given sex) with replacement and by recomputing sliding averages *M*_*j*_, *IQR*_*j*_, and *MC*_*j*_ for each of the 10^4^ samples. The resulting lifetime trajectories 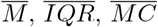, were represented as 20-dimensional vectors (e.g., twenty values 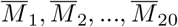 at twenty ages *a*_1_, *a*_2_, …, *a*_20_ ranging from 12 to 70) and collected in two 20 × 10000 dimensional matrices *M*_*female*_ and *M*_*male*_, *IQR*_*f*_ and *IQR*_*m*_, and *MC*_*f*_ and *MC*_*m*_.

The univariate difference between male and female trajectories was assessed with Fisher’s linear discriminant analysis (e.g. [98]). For every matrix pair *X*_*f*_ and *X*_*m*_, we established 20 × 1 dimensional mean vectors *m*_*f*_ and *m*_*m*_, and 20 × 20 dimensional covariance matrices 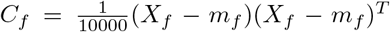 and 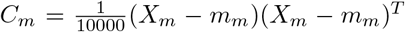, which permitted us to compute the optimally discriminating directions as *w* = (*m*_*f*_ – *m*_*m*_)^*T*^ (*C*_*f*_ + *C*_*m*_)^−1^. After normalizing this 20 × 1 dimensional vector *w*, the projections of all univariate trajectories onto this vector were obtained as *x*_*f*_ = *w*^*T*^ *X*_*f*_ and *x*_*m*_ = *W*^*T*^ *X*_*m*_. Finally, the pairwise distances *x*_*f*_ – *x*_*m*_ between male and female projections were collected into an ‘observed’ distribution.

To test statistical significance, the gender of observers was randomly shuffled 10^4^ times to obtain ‘pseudo-male’ and ‘pseudo-female’ cohorts, and the trajectory vectors were collected into 20×10000 dimensional matrices *X*_*pseudo–f*_ and *X*_*pseudo–m*_. The difference between pseudo-females and pseudo-males was established as described above, and the pairwise distances *x*_*pseudo–f*_ – *x*_*pseudo–m*_ between pseudo-male and pseudo-female projections were collected into a ‘null’ distribution.

Finally, we computed statistical significance (p-value) as the probability that ‘observed’ distances were smaller than ‘null’ distances.

Univariate differences in terms of maturation index MI, principal component PC2, and principal component PC3, were assessed in the same way. Univariate differences in partial trajectories were also assessed in this way, spanning from age 12 to 20, from age 20 to 30, from age 30 to 40, and from age 40 to 70.

### 4.5 Computing maturation index

As all parameters tended to change concomitantly, we sought to summarize the development of all three parameters in terms of a single maturation index. As a first step, averaged distribution parameters 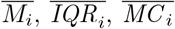 were normalized (z-scored), to obtain values 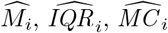 with zero mean and unit variance, over all typically neurotypical observers. Next, we computed the principal component direction over all typical observers (both male and female), which captured most of the variance (~ 80%). We defined the maturation index (MI) as the projection of normalized average parameters 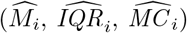 onto this direction, or equivalently, a linear combination of these parameter triplets:

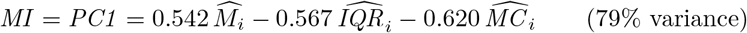

The maturation index provided a convenient summary of binocular rivalry statistics over different ages and sexes. The other components were:

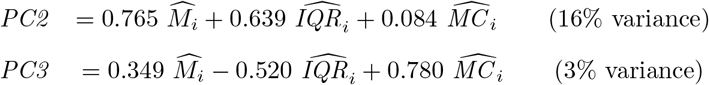

Confidence intervals were computed by repeatedly sampling observers with replacement, and recomputing the average parameters.

### 4.6 Further statistical analysis

Fisher’s linear discriminant analysis (e.g. [98]) was also used to assess multivariate differences between female and male trajectories. Repeated sampling of observers (of a given sex) and recomputing sliding averages resulted in 104 lifetime trajectories, which were represented as 60-dimensional vectors (i.e., value triplets at twenty ages ranging from 12 to 70), and collected in two 60×10000 dimensional matrices *V*_*female*_ and *V*_*male*_. From the mean and covariance of trajectory vectors, we obtained the optimally discriminating axis *w* in 60-dimensional trajectory space, as well as the projections *v*_*f*_ = *w*^*T*^ *V*_*f*_ and *v*_*m*_ = *w*^*T*^ *V*_*m*_ of individually multivariate trajectories onto this axis. The pairwise distances *v*_*f*_ – *v*_*m*_ were collected into an ‘observed’ distribution and compared to the ‘null’ distribution obtained from pseudo-female and pseudo-male trajectories. Multivariate differences in partial trajectories were assessed in analogous ways, spanning from age 12 to 20, from age 20 to 30, from age 30 to 40, and from age 40 to 70.

Multivariate differences in particular trajectory points - starting point (age 12), maturation peak (age 18.5 for females, age 23.5 for males), and end point (age 50) - were analyzed by resampling observers and by projecting trajectory points onto the optimally discriminating axis. Significance of pairwise distances was assessed by comparing the resulting ‘observed’ distribution with zero.

Multivariate differences in the space of model parameters (adaptation strength *ϕ*_*a*_, noise strength *σ*_*n*_, adaptation time constant *τ*_*a*_) or of perceptual parameters (sensitivity, stability, exploration) were also assessed with linear discriminant analysis. The fitting and simulation procedure mapped a set of ages *a*_1_, *a*_2_, … , *a*_*n*_ onto a ‘combined cloud’ in three-dimensional space. After obtaining female and male clouds, as well as the optimally discriminating axis, all points were projected onto this axis and the distribution of pairwise differences was established. Significance was assessed by comparing this distribution to zero.

To assess changes in the slope of the univariate trajectory of principal component PC2, random sampling with replacement (10^4^ samples) was used to obtain partial trajectories before and after the female maturation peak (age 14-18 and age 20-30), before and after the male maturation peak (age 14-24 and age 30-40), during early maturity (age 26 to 40) and during late maturity (from 40 to 50), separately for females and males. For each trajectory segment, a regression line and the sign of its slope was computed. Finally, the pairwise distribution of signs ‘++’, ‘+−’, ‘−+’, ‘−−’) was obtained for successive trajectory segments and the statistical significance of the dominant combination (e.g., ‘+−’) was assessed with a Chi-square test (3 degrees of freedom).

### 4.7 Computational model

Bistable perception was modeled in terms of a dynamic system with competition, adaptation, and noise. The specific formulation we used was introduced by Laing and Chow [53,99,100] and has been analyzed and extended by several other groups [23, 101–104]. Note that the Laing and Chow model describes competitive dynamics within a single receptive field and thus does not account for “mixed” percepts. Modeling mixed states would require multiple models coupled by lateral interactions [105, 106].

The dynamic response *r*_1,2_ of each neural representations is given by

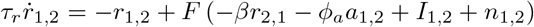

with sensory input *I*_1,2_, intrinsic noise *n*_1,2_, adaptive state *a*_1,2_ and activation function

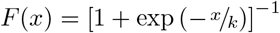

The dynamics of adaptive states *a*_1,2_ is given by

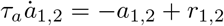

and intrinsic noise *n*_1,2_ is generated from two independent Ornstein–Uhlenbeck processes:

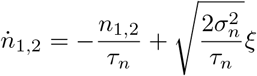

where *ξ* is normally distributed white noise. Different dynamical regimes may be obtained by varying competition (*β*), adaptation (*ϕ*_*a*_, *τ*_*a*_), and noise (*σ*_*n*_), while keeping time constants *τ*_*r*_ and *τ*_*n*_, and activation parameter *k* fixed. Inputs were set as equal (*I*_1_ = *I*_2_ = 1).

### 4.8 Fitting computational model parameters

We performed grid simulations for competition strengths *β* = 1, *β* = 2, *β* = 3, and *β* = 4, because competition strength is not well constrained by our observations. Each simulation lasted 104 s and covered 100-by-100-by-100 value triplets of the other critical model parameters, adaptation strength (*ϕ*_*a*_), time constant of adaptation (*τ*_*a*_), and noise (*σ*_*n*_). The explored range of parameter values was *ϕ*_*a*_ ϵ [0.1, 0.5], *σ*_*n*_ ϵ [0, 0.1] for *β* = 1; *ϕ*_*a*_ ϵ [0.3, 1.2], *σ*_*n*_ ϵ [0, 0.4] for *β* = 2; *ϕ*_*a*_ ϵ [0.5, 2.0], *σ*_*n*_ ϵ [0, 0.4] for *β* = 3; and *ϕ*_*a*_ ϵ [1, 4], *σ*_*n*_ ϵ [0, 0.5] for *β* = 4. For all values of *β*, *τ*_*a*_ was in a range of [0.1, 1.3] s. The remaining, non-critical parameters were *I*_1,2_ = 1, *τ*_*n*_ = 0.1 s, *τ*_*r*_ = 0.02 s, *k* = 0.1, and *dt* = 0.002 s in all four cases.

For each of the 3-by-106 value quadruplets of *β*, *ϕ*_*a*_, *σ*_*n*_, and *τ*_*a*_, we parsed the resulting reversal sequence of 104 s duration into dominance periods by taking *sign*(*r*_1_ – *r*_2_), and calculated three summary statistics of dominance duration: median, interquartile range (IQR), and medcouple (MC).

To compare simulations to the reversal statistics of human observers, we identified combinations of model parameters that reproduce the observed *average* binocular rivalry statistics (median, IQR, MC) of females and males from age 12 to 70. Specifically, we retained all parameter combinations for which the simulated median, IQR and MC fell within 5% of at least one set of observed *average* values.

### 4.9 Simulating experiments with modulated inputs

After fitting model parameter combinations to observers, we performed simulations on these parameter combinations with a time-varying input bias, Δ*I*_*t*_, generated as an Ornstein-Uhlenbeck process with standard deviation *σ*_*I*_ = 0.3 and autocorrelation time *τ*_*I*_ = 0.2 s. To keep total input constant *I*_1_(*t*) + *I*_2_(*t*) = 2, the bias was applied anti-symmetrically *I*_1_(*t*) = 1 + Δ*I*(*t*), *I*_2_(*t*) = 1 − Δ*I*(*t*). The time-scale of this modulation was chosen to obtain a *low* reversal probability, in order to better characterize the circumstances of *failed* reversals.

The simulated dynamics were obtained in time-steps of 1 ms, and included response Δ*r* = *r*_1_ – *r*_2_, perceptual dominance *sign*(Δ*r*), differential adaptation Δ*a* = (*a*_1_ − *a*_2_) *sign*(*r*), and differential input Δ*I* = (*I*_1_ – *I*_2_) *sign*(Δ*r*). Note that positive Δ*a* favours the suppressed percept, whereas positive Δ*I* favours the dominant percept. Intrinsic noise *n*_1,2_ was treated as unobservable, and was subsumed in the probabilistic analysis described in the next section. In other words, probabilities and expectation values were obtained by averaging over intrinsic noise.

### 4.10 Obtaining perceptual parameters

We calculated perceptual parameters from the simulations described above. In the simulated time series, reversals were defined as time-points where responses were equal, *r*_1_ = *r*_2_. To calculate perceptual parameters, we disregarded transition periods, defined as 20 ms before and after a reversal. We classified all other time points as either initiation periods preceding a reversal (40-21 ms before a reversal), or as periods not closely preceding a reversal. Based on this classification, we established the following probabilities: the joint probability of *P* (Δ*I*, Δ*a*), the conditional joint probability, given initiation of a subsequent reversal *P* (Δ*I*, Δ*a* | *init*), and the conditional joint probability of Δ*a* and Δ*I*, given no subsequent reversal, *P* (Δ*I*, Δ*a* | *no rev*). From this, the we calculated conditional probability of a subsequent reversal, given input bias and model response, *P* (*init* | Δ*I*, Δ*a*), which we write *P*_*init*_(Δ*I*, Δ*a*). In the vicinity of the median state (Δ*I*_*M*_, Δ*a*_*M*_), the logarithm of the reversal probability ln *P*_*init*_ varies almost linearly with Δ*I* and Δ*a* (**Fig. 4b**, quality of linear regression *r*^2^ > 0.99).

To characterize the planar surface ln *P*_*init*_(Δ*I*, Δ*a*) in an intuitive and meaningful way, we defined three independent ‘perceptual’ parameters - sensitivity, stability, and exploration. This choice of terms was motivated by the manner in which a dynamical system with two internal states (perception bias Δ*r* and adaptation bias Δ*a*) interacts with a time-varying external state (input bias Δ*I*). The system is less stable for a ‘mismatch’ between internal and external states (Δ*r*, Δ*a* and Δ*I*) and more stable for a ‘match’ between these states. Given a sufficiently large ‘mismatch’, the system responds with a ‘reversal’ (inversion of Δ*r*). Thus, a reversal renders the system more stable by establishing a new ‘match’. Finally, a ‘mismatch’ with internal state Δ*r* can arise both with respect to external state Δ*I* and internal state Δ*a*.

Such an interaction may be characterized in terms of three paramaters. Firstly, the ‘sensitivity’ of the system to ‘mismatch’ of any kind (between Δ*r* on one side and Δ*a* and/or Δ*I* on the other side). Secondly, the degree of ‘stability’ that is typically added by a reversal. Thirdly, the relative sensitivity to internal mismatch Δ*a*, as opposed to external mismatch Δ*I*, which we term ‘exploration’. The precise definitions are given below and are illustrated in **Supplementary Fig. 3**. Auxiliary variables *γ*_*I*_ and *γ*_*A*_ are the partial derivatives *∂*_Δ*I*_ ln *P*_*init*_ ≡ *γ*_*I*_ and *∂*_Δ*a*_ ln *P*_*init*_ ≡ *γ*_*A*_.

In the generally accepted meaning of the term, ‘sensitivity’ describes a proportionality (slope) between stimulus and response. Accordingly, we define ‘sensitivity’ as the derivative of the logarithm of reversal probability (response) with respect to input- and/or adaptation-bias (stimulation). This parameter corresponds to the gradient vector or, equivalently, to the tangent of the slope 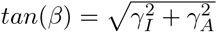 of the planar surface ln *P*_*init*_.

A generally accepted meaning of ‘stability’ is the rareness (inverse probability) of a change. Accordingly, we define ‘ stability’ as the negative logarithm of reversal probability. This corresponds to the negative elevation −*γ*_*I*_ Δ*I*_*M*_ – *γ*_*A*_Δ*a*_*M*_ of the planar surface ln *P*_*init*_ at the typical (median) state (Δ*I*_*M*_, Δ*a*_*M*_) of model and environment after a reversal. A convenient reference level is the value of ln *P*_*init*_ at the unbiased state (Δ*I* = 0, Δ*a* = 0). Thus ‘stability’ reflects the additional degree of stability attained after a reversal, not the absolute level of stability (or instability) imposed by a time-varying environment.

The accepted meaning of ‘exploration’ is the undertaking of risky choices against better knowledge. Here, we appropriate this term for choices against external evidence that are prompted by an ‘internal mismatch’ between Δ*a* and Δ*r*. Specifically, we defined ‘exploration’ as the direction of the gradient vector at the median state 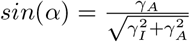, which specifies rotation of the planar surface about the ln *P*_*init*_ axis. While the choice of ‘exploration’ may be less compelling than that of ‘stability’ and ‘sensitivity’, it does describe a third independent aspect of dynamical system behaviour.

## 5 Data availability

The eye tracking datasets are available from the authors on request.

## 6 Computer code

The code used to extract reversal sequences is available via https://github.com/cognitive-biology/Cumulative-smooth- The code used for computational modelling and simulated experiments is available from the authors on request.

## 7 Acknowledgements

We acknowledge support from NKFI (Nemzeti Kutatási, Fejlesztési és Innovációs Hivatal) 110466 and 134370 to IK, and from DFG (Deutsche Forschungsgemeinschaft) BR987/3 and BR987/4 to JB, as well as helpful discussions with Alexander Pastukhov. We thank Péter Soltész, Doctoral School of Psychology, Eötvös Loránd University, Budapest, Hungary for carrying out an extensive pilot study before the current project, and Kinga Farkas, MD, PhD, Department of Psychiatry and Psychotherapy, Semmelweis University, for recruiting and diagnosing patients. We are indebted for the work of Tímea Jéger, a graphic artist who helped us to generate the final versions of the figures.

## 8 Author contributions

GZ designed, developed and performed behavioral experiments; SA analyzed eye movements and conducted computational modelling; ZU recruited and diagnosed BPD patients, all authors discussed the results and contributed to the final manuscript.

## 10 Supplementary figures

**Figure 6: Supplementary figure 1.**
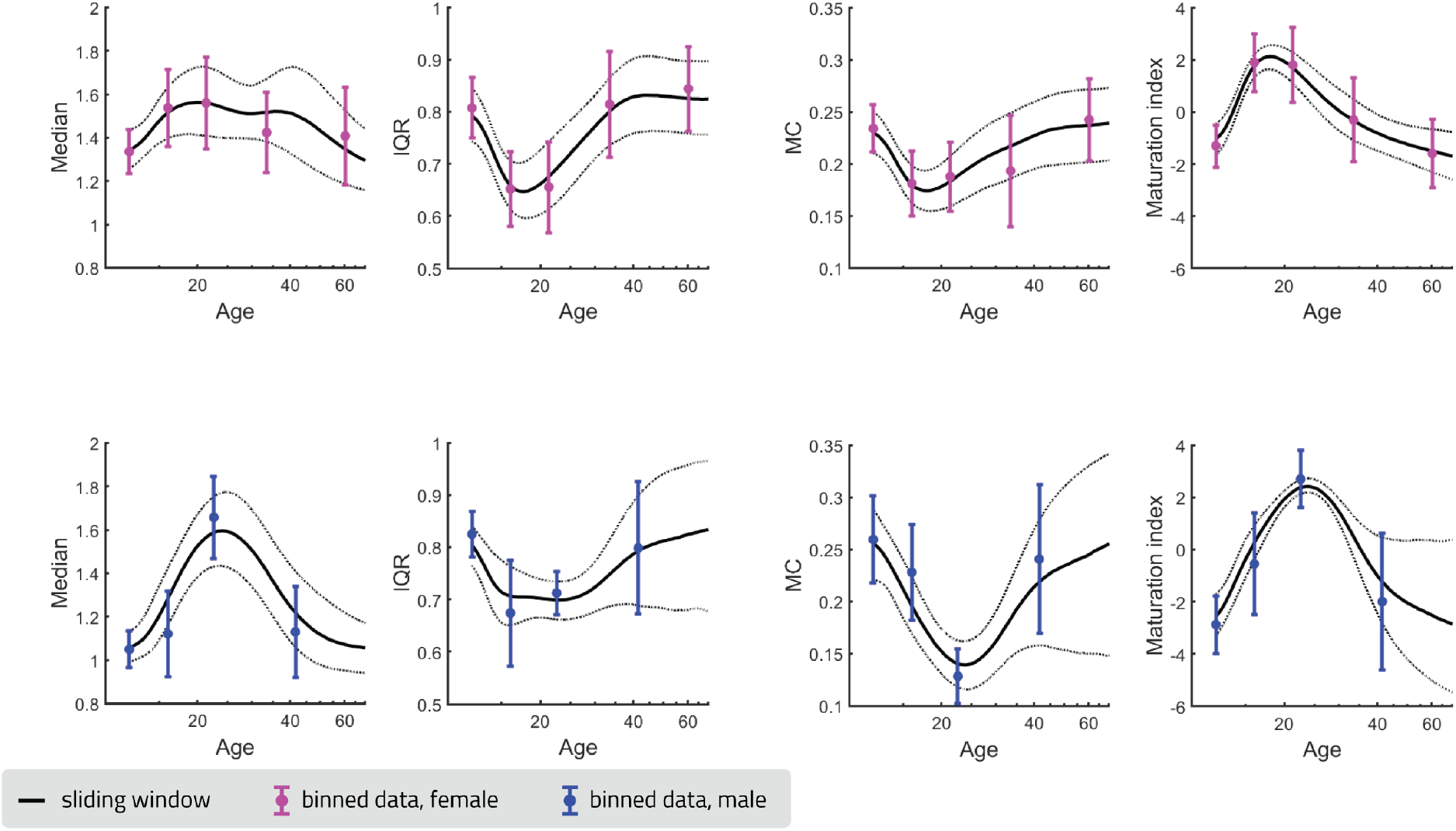
One-dimensional plots of binocular rivalry observables (as in Fig. 2 a, b) and maturation index (as in Fig. 2 d). Symbols and error bars show mean and standard deviation of measured values within non-overlapping age bins. Smooth curves show the median and standard-deviation of Gaussian-weighted sliding-window averages, obtained over random samples of observers (10000 samples with replacement).

**Figure 7: Supplementary figure 2.**
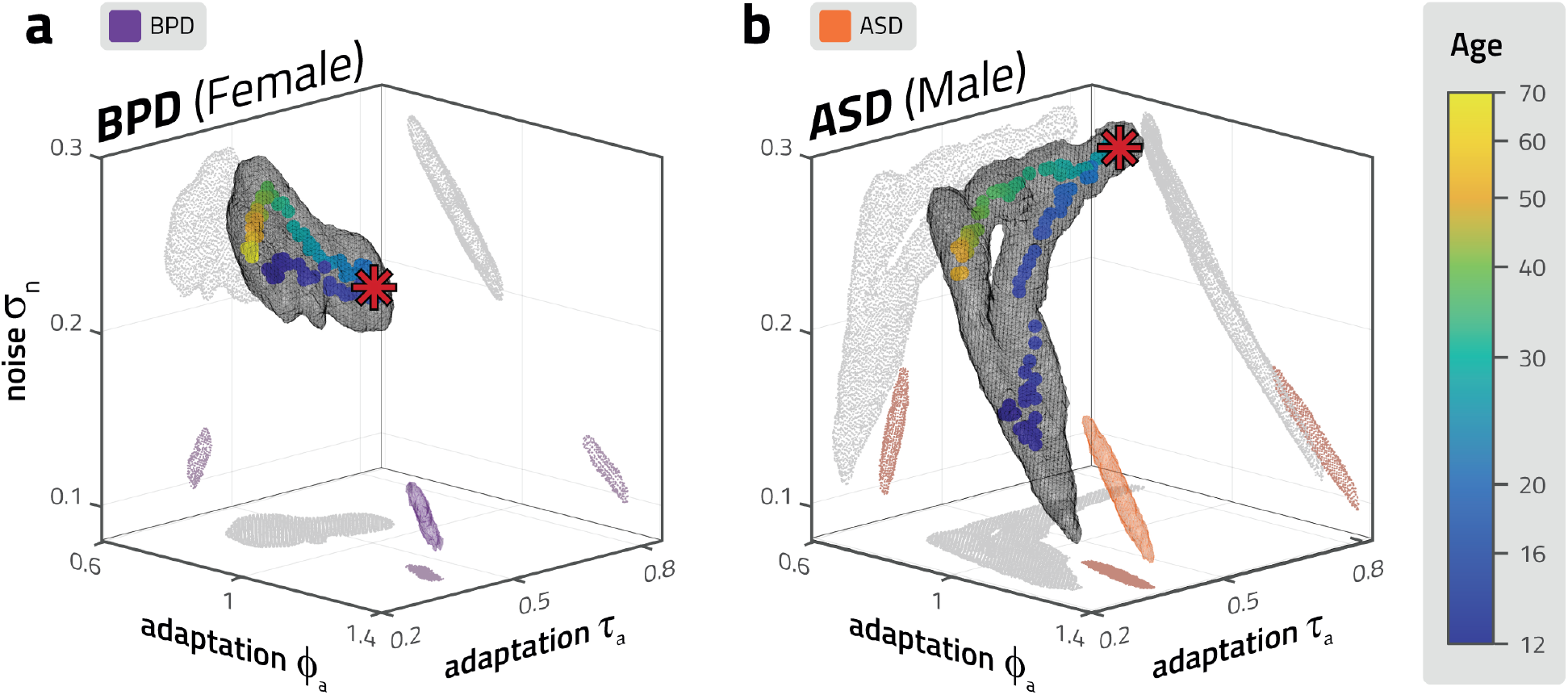
Multistable perception model parameters reproducing observed results of clinical groups. a, b: Combinations of model parameter values reproducing dominance statistics (within a ±5% tolerance range) of clinical observer groups (BPD in purple, ASD in orange) in *ϕ*_*a*_-*τ*_*a*_-*σ*_*n*_ subspace, with constant competition *β* = 3, presented alongside typical developmental and maturational trajectories of the respective sex (same as in Fig 3 c, d).

**Figure 8: Supplementary figure 3.**
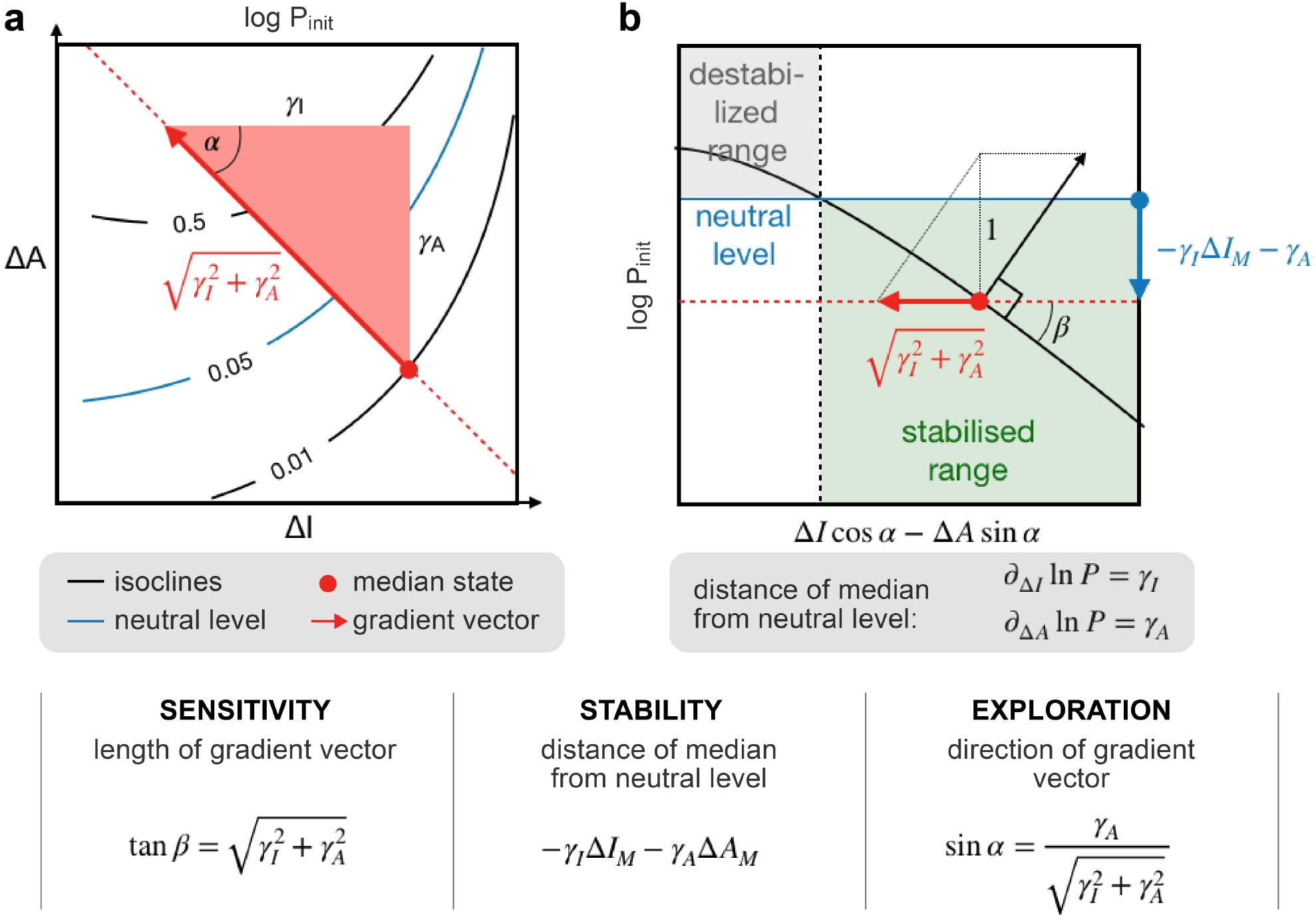
Mathematical definitions of perceptual parameters. a: The conditional likelihood of reversals as a function of Δ*I* and Δ*a* values during initiation periods, in the vicinity of the median state (over all periods, red dot). The reversal probability grows exponentially to the direction of the red line. Sensitivity is defined as the length of the gradient vector *∂* ln *P*_*init*_(Δ*I*, Δ*a*).Exploration is defined as the direction of the gradient vector, *α*. b: Enlarged cut through the ln *P* (Δ*I*, Δ*a*) surface in the direction of the the gradient vector (red arrow), showing the neutral level (defined by Δ*I* = Δ*a* = 0, black dashed line) and median level (red dashed line). Stability is defined as the distance between these levels. At positive values of stability (stabilized range), median states lower reversal probability; vice versa for negative values (destabilized range). Auxiliary variables *γ*_*I*_ and *γ*_*A*_ are the partial derivatives *∂*_Δ*I*_ ln *P*_*init*_ ≡ *γ*_*I*_ and *∂*_Δ*a*_ ln *P*_*init*_ ≡ *γ*_*A*_.

